# CD8^+^ T cell priming in oral mucosa-draining lymph nodes supports systemic immunity

**DOI:** 10.1101/2020.09.21.306449

**Authors:** Juliana Barreto de Albuquerque, Diego von Werdt, David Francisco, Geert van Geest, Lukas M. Altenburger, Jun Abe, Xenia Ficht, Matteo Iannacone, Remy Bruggmann, Christoph Mueller, Jens V. Stein

## Abstract

The gastrointestinal (GI) tract constitutes an essential barrier against ingested pathogens. While immune reactions are well-studied in the lower GI tract, it remains unclear how adaptive immune responses are initiated during microbial challenge of the oral mucosa, the primary site of pathogen encounter in the upper GI tract. Here, we identify mandibular lymph nodes (mandLN) as sentinel lymphoid organs that collect orally administered *Listeria monocytogenes* (Lm), leading to local CD8^+^ T cell activation. In contrast to CD8^+^ T effector cells (T_EFF_) generated in mesenteric lymph nodes, mandLN CD8^+^ T_EFF_ lacked a gut-seeking phenotype but contributed to systemic host protection. Accordingly, mandLN stromal and dendritic cells expressed low levels of enzymes required for gut homing imprinting. Our findings extend the concept of regional specialization of immune responses along the length of the GI tract, with mandLN acting as oral lymph-draining counterparts of intestinal lymph-draining LN of the lower GI tract.

**Summary:** *Listeria monocytogenes* ingestion leads to priming of cytotoxic T cells in oral mucosa draining mandibular lymph nodes, which contribute to systemic host protection.

## Introduction

The digestive, or gastrointestinal (GI), tract constitutes the major external surface of the human body and comprises the upper GI tract with oral cavity, pharynx and esophagus, and the lower GI tract with stomach, small and large intestine and rectum. The GI tract needs to offset permissive uptake and digestion of nutrient and water with protection against invading pathogens. Accordingly, GI immune responses range from tolerance against commensals and food antigens to reactivity against ingested pathogens. Innate and adaptive immune responses have been most extensively studied in the lower GI tract (Kiyono and Azegami, 2015; Faria et al., 2017; Belkaid and Harrison, 2017; Schulz and Pabst, 2013; Shale et al., 2013; Mowat and Agace, 2014). Gut-associated lymphoid tissues (GALT) of the small intestine such as Peyer’s patches (PP) form together with gut lymph nodes (gLN) complementary inductive sites for intestinal immune reactions (Brandtzaeg et al., 2008). M cells embedded in the epithelium overlying GALT sample luminal Ag, while gLN including mesenteric LN (MLN) screen intestinal lymph to intercept pathogens, which have breached the epithelial barrier. Both MLN and PP promote the generation of gut-homing α4β7^high^ CCR9^+^ CD4^+^ and CD8^+^ T effector cells (T_EFF_) (Mora et al., 2003). This phenomenon relies on retinoic acid (RA) generation by the retinal aldehyde dehydrogenase (RALDH)-expressing CD103^+^ migratory dendritic cells (DC) and stromal cells of intestinal Ag-sampling lymphoid tissue, which induces a gut-seeking phenotype in activated lymphocytes (Iwata et al., 2004; Erkelens and Mebius, 2017; Larange and Cheroutre, 2016). Recent findings have further refined our understanding of gut immunity by identifying compartmentalized tolerogenic and inflammatory CD4^+^ T cell immune responses in individual gLN, which drain lymph from proximal versus distal segments of the intestinal GI tract (Esterhazy et al., 2019). Thus, the lower GI tract is characterized by regional specialization of adaptive immune reactions according to the local microenvironment (Mowat and Agace, 2014).

The upper GI tract, in particular the mucosa of the oral cavity, also contains an abundant microbiota and constitutes the first site of contact with dietary Ag and ingested pathogens (Moutsopoulos and Konkel, 2018; Gaffen and Moutsopoulos, 2020). The oral barrier has recently gained attention for its complex immune network characterized by abundant Th17 CD4^+^ T cells and numerous macrophage subsets (Park et al., 2017; Moutsopoulos and Konkel, 2018; Gaffen and Moutsopoulos, 2020). In contrast, the initiation of adaptive immune responses against microbes sampled from oral mucosa has not been well characterized, in particular for cytotoxic CD8^+^ T cell responses that occur in response to intracellular pathogens of the oral cavity. In this context, mice lack tonsils, and lumen-sampling M cells are restricted to nasopharynx-associated lymphoid tissue (NALT) of the nasal cavity (Pabst, 2015). Yet, to the best of our knowledge, no comprehensive attempt has been made to map oral mucosa-draining LN, although in a rodent periodontitis model, antibacterial responses are detectable in mandibular and accessory mandibular LN (here collectively abbreviated as mandLN; also sometimes referred to as superficial cervical LN) (Mkonyi et al., 2012; Van den Broeck et al., 2006; Lohrberg and Wilting, 2016). Our current understanding is limited since most oral immunization models in small rodents bypass the oral cavity by employing intragastric (i.g.) gavage. In models where pathogens are administered intraorally (i.o.), the analysis of immune responses often excludes lymphoid tissue of the neck and head region (D’Orazio, 2014; Sheridan et al., 2014). Taken together, it remains unclear how microbial uptake by the oral mucosa initiates local priming and whether this process impacts on distal immune responses (Moutsopoulos and Konkel, 2018).

Here, we administered *Listeria monocytogenes* (Lm) into the oral cavity of immunocompetent mice to mimic a first encounter with an ingested microbe. Lm is a food-borne gram-positive bacterium, which infects macrophages and hepatocytes in its target organs spleen and liver and is cleared by cytotoxic effector CD8^+^ T_EFF_. We uncovered early priming and proliferation of Ag-specific CD8^+^ T cells in mandLN that preceded activation kinetics in spleen, MLN and PP. In contrast to CD8^+^ T_EFF_ primed in MLN after i.g. Lm gavage, mandLN CD8^+^ T_EFF_ did not display a gut-homing α4β7^high^ CCR9^+^ phenotype. Instead, mandLN CD8^+^ T_EFF_ contributed to systemic host protection similar to T_EFF_ generated in spleen by i.v. Lm infection. This finding correlated with low expression of genes imprinting gut homing in the mandLN microenvironment. Our observations expand the emerging concept of a compartmentalized host immune response along the length of the entire GI tract according to the Ag sampling location, leading to preferential generation of systemically disseminating T_EFF_ in oral mucosa-draining mandLN and gut-seeking T_EFF_ in MLN.

## Results and discussion

### Ingested Lm is intercepted in mandLN and leads to local LN hyperplasia

To examine drainage of oral mucosa lymph, we injected a lymph tracer into gingiva and observed rapid accumulation (≤ 5 min) in draining mandLN (**Fig. 1A**), suggesting a connecting lymphatic network between oral cavity and mandLN. In tissue sections, the gingiva adjacent to teeth was reported to contain numerous lymphatic vessels, which are located more superficially than in most mucosal tissues (Ushijima et al., 2008; Mkonyi et al., 2012). To obtain an overview of the non-sectioned oral lymphatic network, we carefully exposed the mandibular gingiva and draining mandLN in a Prox1-GFP reporter mouse strain for lymphatic vessels (Choi et al., 2011) (**Fig. S1A**). We confirmed the presence of an extensive gingival lymphatic network around the mandibular incisors (**Fig. 1B, C**) connected to mandLN (**Fig. 1D**).

**Figure 1.**
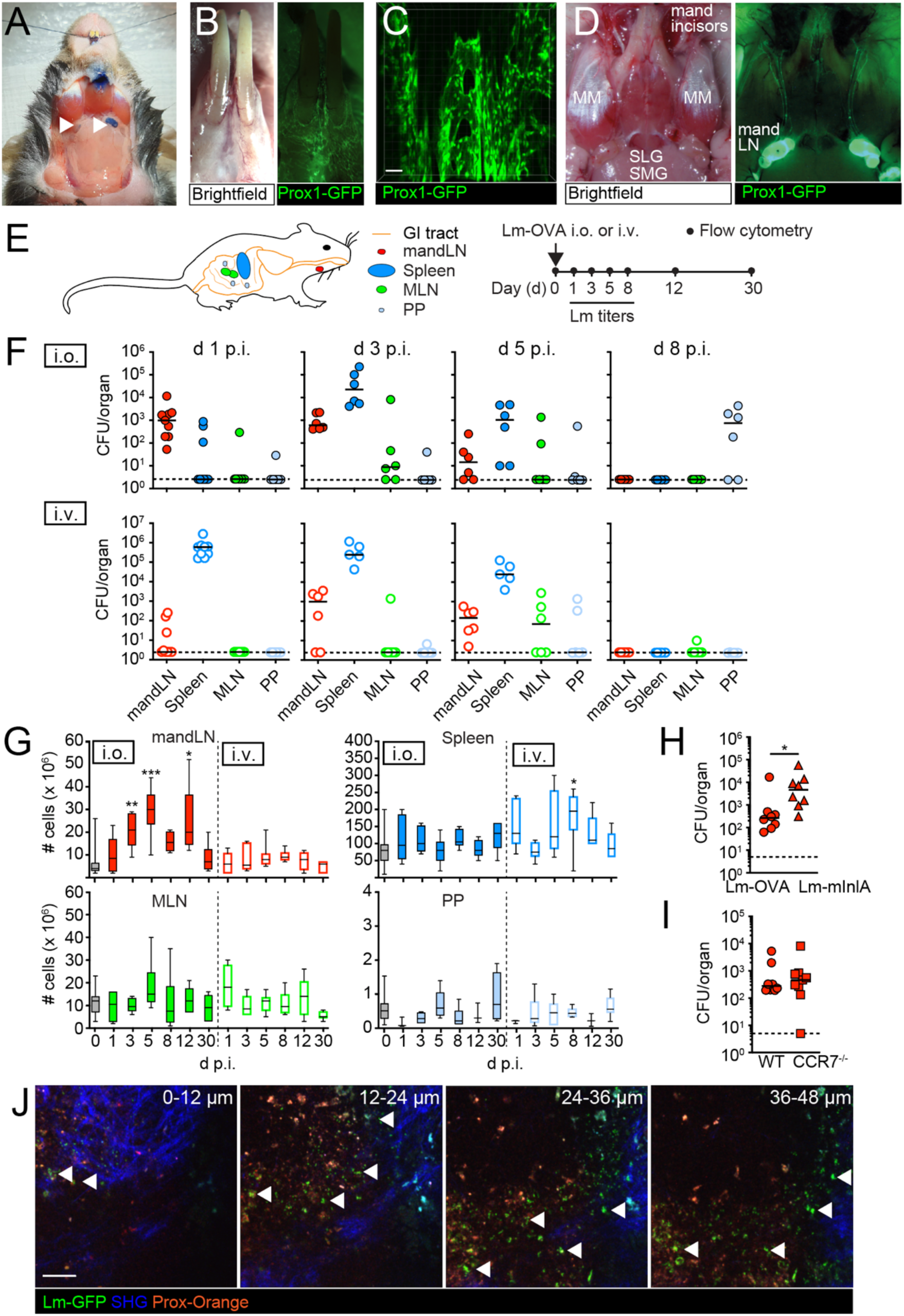
Lm inoculation into oral cavity leads to rapid bacterial accumulation in mandLN. **A**. Drainage of a tracer injected into the gingiva to mandLN (arrowheads). **B**. Overview of lymphatic vessel network in gingiva of Prox1-GFP reporter mouse. **C**. 2PM reconstruction of lymphatic network in gingiva surrounding mandibular incisors. Scale bar, 20 µm. **D**. Overview of lymphatic vessels between mandibular incisors and mandLN of Prox1-GFP reporter mouse. MM, masseter muscle; mandLN, mandibular lymph node; SLG, sublingual salivary gland; SMG, submandibular salivary gland. **E**. Experimental setup for oral infection. **F**. Lm-OVA CFU per lymphoid organ after intraoral (i.o.) *versus* intravenous (i.v.) Lm infection. Lines depict median, dotted line is limit of detection. **G**. Lymphoid organ cellularity during i.o. and i.v. Lm-OVA infections. **H**. CFU counts in mandLN after i.o. infection of Lm-OVA *versus* Lm-InlA^m^. **I**. Lm-OVA CFU counts in mandLN after i.o. infection of WT or CCR7^-/-^ mice. **J**. Lm-GFP accumulation (arrowheads) after i.o. infection at distinct depths from mandLN capsule. SHG, second harmonic generation. Scale bar, 40 µm. Data in F and G are from 2-3 independent experiments with each 2-3 mice/time point. Data in G were analyzed using a Kruskal-Wallis test against “d 0”. Data in H and I are pooled from 2-3 independent experiments with each 4-5 mice/group and analyzed using a Mann-Whitney test. *, p < 0.05; **, p < 0.01; ***, p < 0.001.

We investigated the relevance of lymphatic drainage in a model of oral infection. Lm expressing the model antigen ovalbumin (Lm-OVA) (Zehn et al., 2009) was directly administered into the oral cavity of C57BL/6 mice. In the first 8 days post i.o. infection, we isolated mandLN, spleen, MLN and PP of infected mice and determined Lm-OVA spread by colony forming unit (CFU) analysis. We compared these values to CFU counts after i.v. administration of Lm-OVA (**Fig. 1E**). We observed that mandLN contained the highest Lm-OVA CFU per organ on d 1 following i.o. infection, whereas spleen contained the highest CFU following systemic administration (**Fig. 1F**). On d 3 after i.o. infection, Lm-OVA became also detectable in spleen and MLN (**Fig. 1F**). Nonetheless, Lm-OVA CFU counts, when detectable, remained lower for MLN than mandLN and were barely detectable in PP before d 8 after i.o. infection (**Fig. 1F**). This is in line with the notion that > 90% of ingested bacteria are killed by gastric acids (Saklani-Jusforgues et al., 2000; Pitts and D’Orazio, 2018). Residual Lm in the intestinal lumen may have been degraded by digestive enzymes, or failed to cross the mucus layer of the intestinal epithelium or their tight junctions, or become cleared by peristaltic contraction and mucus secretion. Together, these factors likely contribute to the thwarted and delayed onset of Lm capture in PP and MLN in our model of oral infection.

To explore whether Lm retention leads to increased lymphocyte numbers in reactive LN, we determined lymphoid organ cellularity following i.v. and i.o. Lm-OVA infection. We observed a rapid onset of mandLN hyperplasia following i.o. infection, consistent with local bacterial sampling at early time points (**Fig. 1G**). In contrast, i.v. Lm-OVA infection did not cause mandLN hypercellularity at any time point analyzed, whereas splenocyte numbers became significantly increased on d 8 p.i. (**Fig. 1G**). In our model of oral infection, Lm-OVA did not cause significant increases in MLN and PP cellularity, reflecting limited CFU detection in the first week p.i. at these sites (**Fig. 1G**).

Expression of a mutant internalin InlA^m^ increases its binding affinity to mouse E-cadherin by four orders of magnitude (Wollert et al., 2007). As a result, i.g. gavage of Lm-InlA^m^ leads to a higher *in vivo* virulence as compared to non-murinized InlA Lm strains (Monk et al., 2010). To examine whether the interplay of InlA and E-cadherin facilitates early Lm accumulation in mandLN, we compared CFU counts of Lm-OVA and Lm-InlA^m^ after i.o. infection. Lm-InlA^m^ CFU counts in mandLN were significantly increased at 24 h p.i. as compared to Lm-OVA, suggesting that E-cadherin binding to oral mucosa epithelium might increase transport to mandLN (**Fig. 1H**). As caveat, this finding may have been influenced by the higher virulence of the Lm strain carrying the InlA mutation as compared to the Lm-OVA strain (Bécavin et al., 2014). To further investigate the mechanism underlying the rapid appearance of ingested Lm in mandLN, we i.o. administered Lm-OVA in CCR7^-/-^ recipients, which lack DC trafficking from peripheral tissues to sentinel LN (Förster et al., 2008; Schulz et al., 2016).

Bacterial loads in CCR7^-/-^ mandLN on d 1 p.i were comparable to WT values, suggesting that active DC migration to mandLN was not a limiting factor for initial Lm accumulation in draining LN (**Fig. 1I**). These observations prompted us to examine a passive transport mechanism for ingested Lm via oral cavity-draining lymphatic vessels. We administered GFP-expressing Lm (Lm-GFP) onto the gingiva of mice prepared for intravital imaging to directly explore its retention in draining mandLN. To this end, we adapted a model originally developed for submandibular salivary gland surgery (Ficht et al., 2018) to visualize the adjacent mandLN (**Fig. S1B**). Lm-GFP accumulated in the subcapsular sinus (SCS) of mandLN within 120 min post oral deposition, as assessed by two-photon microscopy (2PM)-based intravital imaging (**Fig. 1J**). Taken together, these data suggest that oral uptake of Lm leads to its drainage to mandLN, at least in part via afferent lymphatics of the oral mucosa.

### Ag-specific CD8^+^ T cells form clusters and dynamically interact with CD11c^+^ cells mandLN following i.o. Lm infection

To quantify the relevance of Lm capture in mandLN on CD8^+^ T cell activation, we transferred fluorescent protein-expressing OT-I TCR tg T cells, which recognize the OVA_257-264_ peptide in the context of H-2K^b^ (Hogquist et al., 1994), together with fluorescently labeled polyclonal CD8^+^ T cells into CD11c-YFP hosts. In these recipients, CD11c^+^ cells including antigen-presenting DC express YFP (Lindquist et al., 2004). One day later, we i.o. infected mice with Lm-OVA and isolated mandLN at 1, 2 and 3 d p.i. for histological analysis. At 1 d p.i., we observed occasional large clusters of OT-I but not polyclonal CD8^+^ T cells around CD11c^+^ cells in both mandibular and accessory mandibular LN, suggesting early Ag-driven interactions (**Fig. 2A**, left panel). These clusters became smaller on d 2 and 3 post i.o. infection (**Fig. 2A**, middle and right panel). As changes in dynamic T cell behavior act as sensitive indicator for priming and precede detectable expression of activation markers (Moreau and Bousso, 2014; Sivapatham et al., 2019), we performed 2PM of mandLN in C57BL/6 or CD11c-YFP recipients containing tdT^+^ or dsRed^+^ OT-I T cells and polyclonal control CD8^+^ T cells during the first 72 h post i.o. Lm-OVA infection. To benchmark physiological T cell behavior in steady state, we analyzed naïve OT-I motility parameters in mandLN of uninfected mice. OT-I T cells displayed high speeds (15.4 ± 5.1 µm/min; mean ± SD), low arrest coefficients (median of 1.6% of track segments < 5 µm/min; **Fig. 2C and D**) and a corrected track straightness of 8.3 ± 2.7 (median ± SD, corresponding to a non-corrected meandering index of 0.61 ± 0.21) (Beltman et al., 2009) (**Video S1**). These values are comparable to T cell motility parameters observed in resting skin-draining LN (Germain et al., 2006; Mempel et al., 2004; Breart and Bousso, 2006), and serve as reference for inflammation-induced changes of dynamic T cell behavior. At d 1 post i.o. Lm infection, polyclonal CD8^+^ T cells displayed a random motility pattern comparable to T cell migration in non-inflamed mandLN, albeit with slightly decreased speeds (12.0 ± 4.5 µm/min) and increased arrest coefficients (**Fig. 2C and 2D**). This finding is in line with mild motility changes displayed by non-cognate T cells in reactive LN (Mempel et al., 2004). In contrast, many OT-I T cells were found to cluster around CD11c^+^ APC on d 1 and 2 p.i. (**Fig. 2B; Video S2**), and most OT-I T cells, in particular in clusters, displayed decreased speeds and high arrest coefficients (**Fig. 2C and 2D**). Such a behavior is consistent with interaction dynamics driven by high cognate pMHC levels displayed on APC (Henrickson et al., 2008; Sivapatham et al., 2019). Starting on d 2 post i.o. Lm infection, OT-I T cell speeds and arrest coefficients began to recover, suggesting decreasing Lm-derived Ag presentation (**Fig. 2C and D**). On d 3 post i.o. Lm infection, OT-I cells ceased to cluster and showed speeds and arrest coefficients largely comparable to polyclonal CD8^+^ T cells (**Fig. 2C and D; video S3**). The analysis of the corrected track straightness followed a similar pattern, with OT-I T cells showing transient confinement on d 1 and 2 p.i., which became less apparent on d 3 p.i. (**Fig. 2E**). Taken together, our confocal and dynamic imaging data are consistent with fast and efficient antigen processing in oral mucosa-draining mandLN, leading to cognate T cell – APC interactions within the first 2 d post i.o. infection.

**Figure 2.**
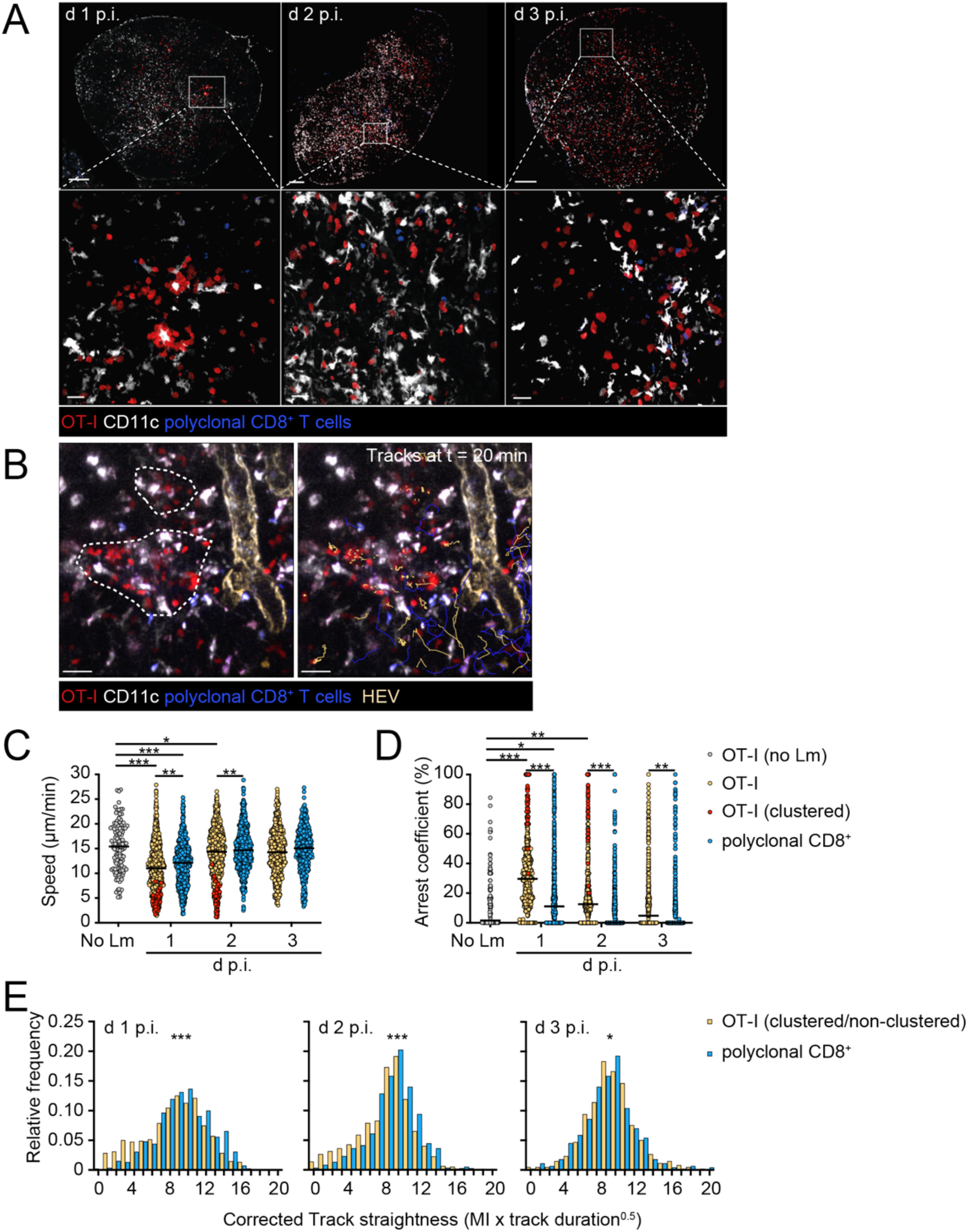
Visualization of CD8^+^ T cell dynamics in mandLN after oral Lm uptake. **A**. Confocal images of mandLN sections after i.o. Lm-OVA administration. Scale bar, 200 (upper) and 20 (lower) µm. **B**. Representative 2PM images of infected mandLN of CD11c-YFP host containing OT-I cells and polyclonal T cells at d 2 p.i.. Left panel shows OT-I clusters around CD11^+^ cells (dotted line), right panel shows overlaid tracks of OT-I (yellow) and polyclonal CD8^+^ T cells (blue). Scale bar, 20 µm. **C-E**. Quantification of imaging data. Track speeds (**C**), arrest coefficient (**D**), and corrected track straightness (**E**). Lines in C and D depict median. Data in C were analyzed using ANOVA with Sidak’s multiple comparison test, in D using Kruskal-Wallis with Dunn’s multiple comparison test and in E using Mann-Whitney test. Data are representative (A) or pooled (C-E) from at least two independent experiments. *, p < 0.05; **, p < 0.01; ***, p < 0.001.

### Oral infection leads to rapid CD8^+^ T cell activation in mandLN over a wide range of Lm inocula

The imaging data suggested cognate OT-I T cell priming by CD11c^+^ antigen-presenting DC. We therefore compared the activation status of migratory and resident DC in resting and i.o. challenged mandLN by flow cytometry 24 h after i.o. Lm-OVA infection. Both populations showed higher and/or more frequent expression of CD80 and CD86 as compared to DC isolated from non-infected mandLN, in line with an activated phenotype (**Fig. S2A-C**). In particular, the proportion of CD86-expressing CD11c^high^ MHC II^+^ resident DC increased from 29 to 47%, with an MFI increase of 35% (mean of two independent experiments with n = 8 control and 8 Lm-infected mice).

We correlated these observations with induction of early activation markers CD69 and CD25 on OT-I T cells following i.o. Lm-OVA infection (for gating strategy see **Fig. S2D**). One d post i.o. Lm infection, mandLN OT-I T cells displayed increased levels of CD69, followed by augmented CD25 expression starting on d 2 p.i. (**Fig. 3A and 3B**). At this time point, OT-I T cells in spleen and MLN showed a delayed and less pronounced increase in these markers compared to mandLN OT-T cells (**Fig. 3A and 3B**). Similar results were obtained for PP OT-I T cells (not shown).

**Figure 3.**
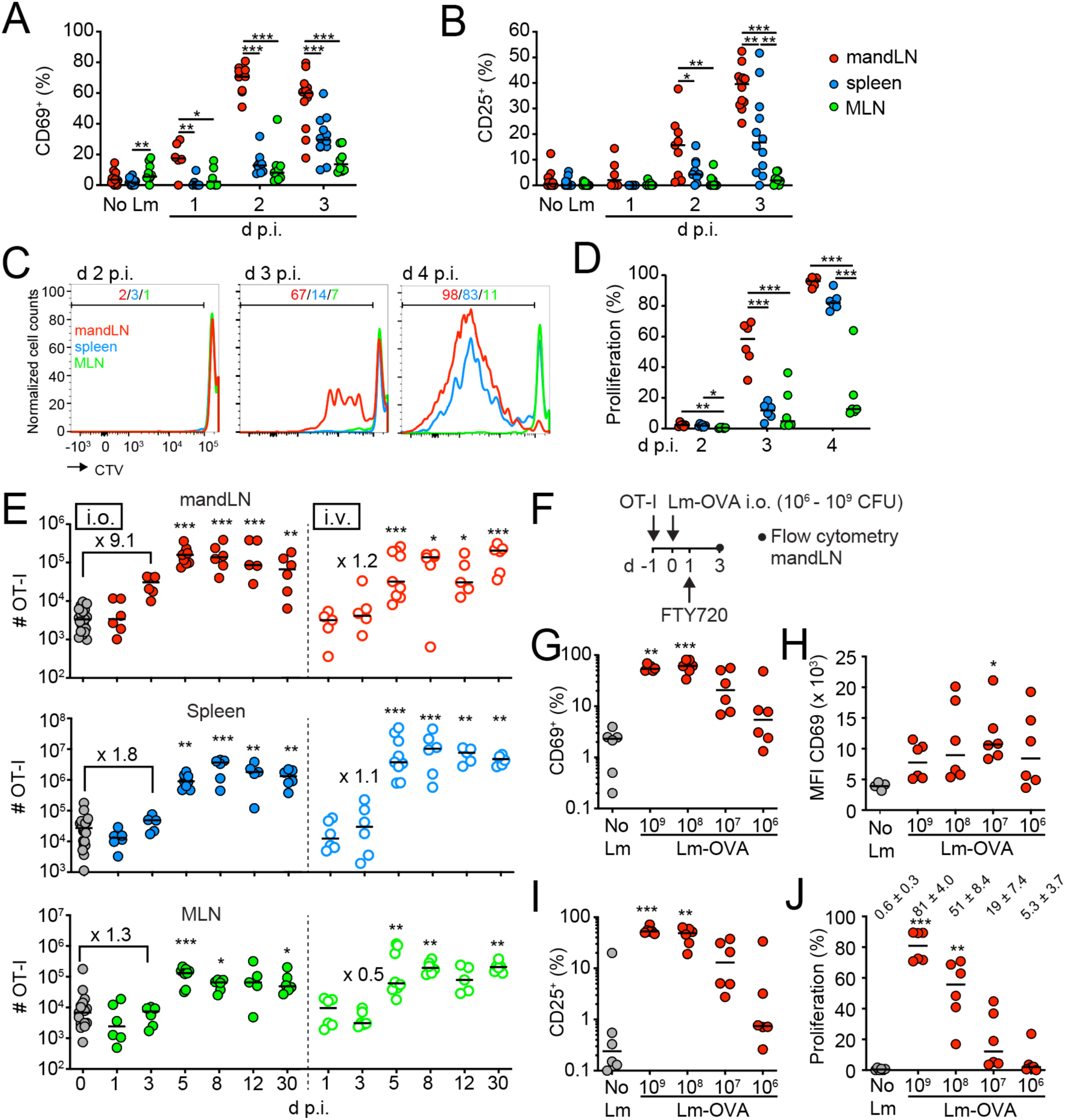
Oral Lm uptake triggers rapid CD8^+^ T cell activation and proliferation in mandLN. **A and B**. CD69 (**A**) and CD25 (**B**) expression in OT-I cells isolated from mandLN, spleen and MLN after i.o. Lm-OVA infection. **C**. Representative flow cytometry plots of OT-I T cell proliferation after i.o. Lm-OVA infection. Numbers indicate percent proliferated cells. **D**. OT-I proliferation in mandLN, spleen and MLN after i.o. Lm infection. **E**. OT-I cell number in lymphoid organs after i.o. and i.v. infection. Numbers depict fold increase of median in d 3 OT-I cell numbers over d 0. **F**. Experimental layout for Lm-OVA inoculum titration. **G-J**. CD69 expression (**G**), MFI CD69 (**H**), CD25 expression (**I**) and proliferation (**J**) on mandLN OT-I T cells after titrating Lm-OVA inoculum. Numbers in J indicate median ± SEM. Data in A and B are pooled from 2-5 experiments per time point with 3 mice per experiment. Data in D are pooled from 2 independent experiments. Data in E and G are pooled from 2-3 independent experiments with each 2-3 mice/group/time point and analyzed using a Kruskal-Wallis test against “d 0” and “no Lm”, respectively. Lines in A, B, D, E and G-J depict median. *, p < 0.05; **, p < 0.01; ***, p < 0.001.

Next, we examined the onset of cell proliferation. To restrict potential cross-contamination by interorgan cell trafficking after Lm infection, we performed these experiments in the presence of FTY720, which sequesters T cells in lymphoid tissue. In line with increased CD25 expression, more than half (58 ± 14.7%; median ± SD) of mandLN OT-I T cells had undergone cell proliferation at d 3 post i.o. infection *versus* 12 ± 5.3% and 5 ± 14% of spleen and MLN OT-I T cells, respectively (**Fig. 3C and 3D**). This trend continued d 4 post i.o. infection (**Fig. 3D**) and was reflected by an earlier and more pronounced expansion of OT-I T cells in mandLN after i.o. *versus* i.v. in the first 5 d p.i. (**Fig. 3E**).

Irrespective of the route of Lm infection, mandLN T_EFF_ were CD44^high^ CD62L^high or low^ on d 5 p.i., whereas spleen T_EFF_ were predominantly CD44^high^ CD62L^low^ (**Fig. S2E and S2F**). In the memory phase (d 30 p.i.), most mandLN OT-I T cells showed a CD44^high^ CD62L^high^ central memory-like phenotype while spleen also contained a minor population of CD44^high^ CD62L^low^ effector memory-like cells (**Fig. S2F**), again independent of the route of infection. We further examined OT-I differentiation into CD127^-^ KLRG-1^+^ short-lived effector cells (SLEC) and CD127^+^ KLRG-1^-^ memory precursor effector cells (MPEC) after oral *versus* systemic Lm infection (**Fig. S2E**). In both routes of infection, we observed comparable SLEC and MPEC proportions on d 5 p.i. (**Fig. S2G**). At d 30 p.i., KLRG1^-^ CD127^+^ central memory-like T cells prevailed in mandLN and spleen after either infection route (**Fig. S2G**).

To test the impact of initial bacterial load on local OT-I T cell activation, we titrated the Lm inoculum and measured OT-I T cell activation and proliferation on d 3 post i.o. infection (**Fig. 3F; Fig. S2H**). We observed a dose-dependent effect on total mandLN cellularity and percent CD69^+^ endogenous CD8^+^ T cells, which became less pronounced with decreasing Lm inoculum (**Fig. S2I and S2J**). This was reflected by a Lm inoculum-dependent activation marker expression and proliferation in OT-I T cells (**Fig. 3G-J**). Notably, we observed increased expression of CD69 and CD25, as well as proliferation with an inoculum of only 10^6^ Lm (**Fig. 3G-J; Fig. S2H**). In sum, our data show that early T cell interactions with CD11c^+^ antigen-presenting cells accelerates T_EFF_ generation in oral mucosa-draining mandLN. Furthermore, we observed that across four orders of magnitude of initial Lm inoculum, sufficient antigenic material is collected by sentinel mandLN to induce detectable CD8^+^ T cell responses.

### MandLN T_EFF_ contribute to the early peripheral cytotoxic T cell pool in spleen

To quantify to which extent mandLN OT-I cells contribute to the early peripheral CD8^+^ T_EFF_ pool, we performed i.o. infections in absence or presence of FTY720 to prevent egress from the priming lymphoid organ (**Fig. 4A**). As predicted, FTY720 treatment led to a 4.4x- and 5x-fold increase in mandLN OT-I T_EFF_ on d 4 and 5 post i.o. Lm infection, respectively. This increase was mirrored by a concomitant decrease in spleen OT-I T_EFF_ from 125 ± 39 to 41 ± 7 x 10^4^ cells/organ (mean ± SEM) in the absence and presence of FTY720, respectively, on d 4 p.i., corresponding to a 67% reduction (**Fig. 4B**). The decrease in splenic OT-I T_EFF_ numbers became a non-significant tendency on d 5 post i.o. infection (499 ± 165 and 209 ± 54 x 10^4^ cells/organ in absence and presence of FTY720, respectively; **Fig. 4B**). The partial rescue of spleen OT-I numbers on d 5 likely reflects Lm spread to spleen by d 3 p.i. and onset of OT-I cell division by d 4 p.i. (**Fig. 1B and Fig. 3D**). In contrast, MLN OT-I T cell numbers on d 4 or 5 p.i. were not altered by FTY720 treatment (**Fig. 4B**). These data suggest that initial CD8^+^ T cell activation in mandLN leads to a rapid release of OT-I T_EFF_ into the circulation, which in part relocate to spleen and represent a sizeable proportion of circulating cytotoxic T cells early after infection.

**Figure 4.**
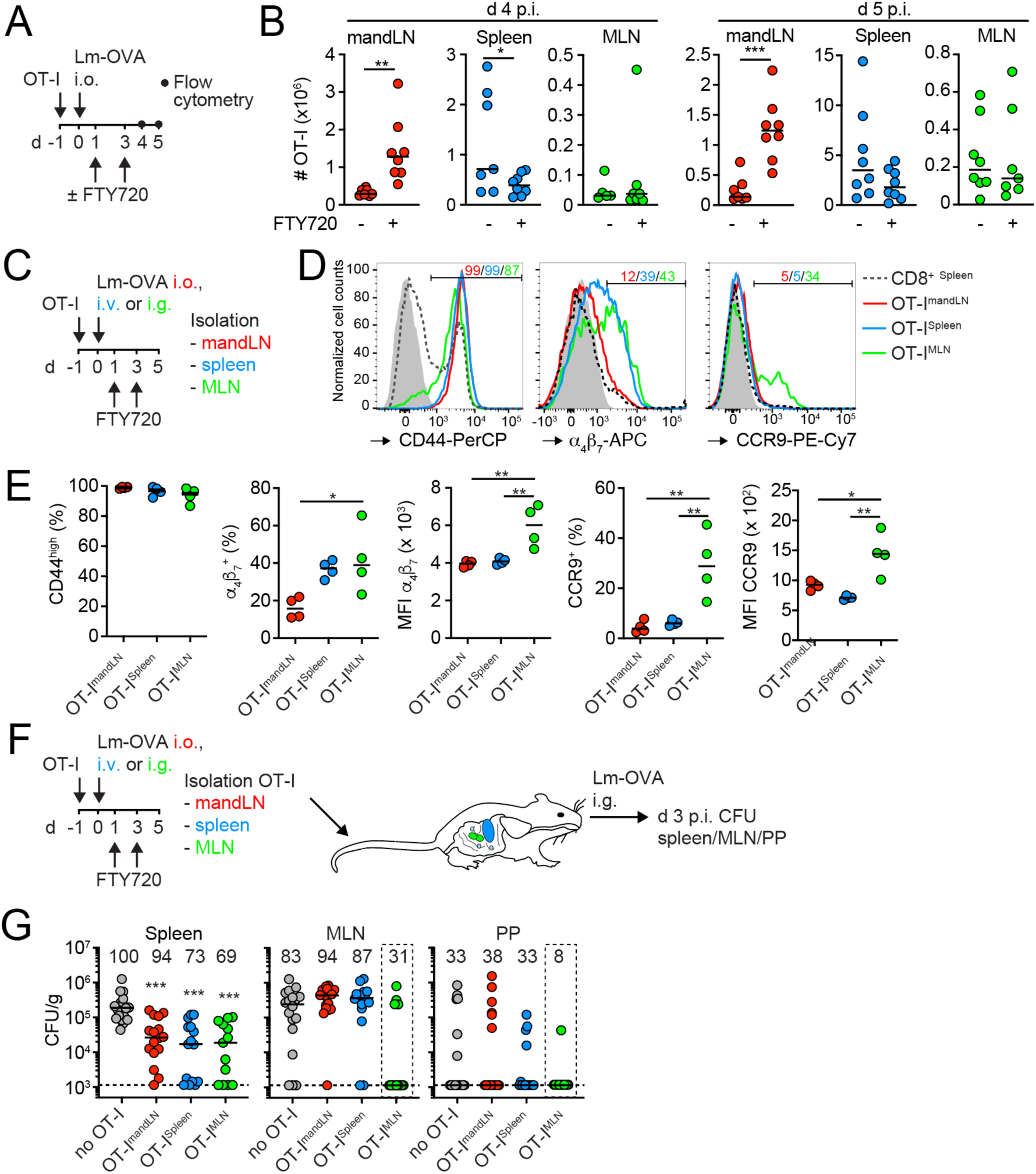
mandLN effector CD8^+^ T cells contribute to systemic immunity. **A**. Experimental layout of egress blockade. **B**. OT-I cell numbers in mandLN, spleen and MLN on d 4 and 5 post i.o. Lm-OVA infection in presence or absence of FTY720. **C**. Experimental layout of homing receptor analysis. **D**. Representative flow cytometry plots of CD44, α4β7 and CCR9 expression in OT-I T_EFF_ on day 5 post Lm-OVA infection via i.o., i.g. and i.v. route. Gray line, FMO control. Numbers depict percent positive cells. **E**. Quantification of percent positive and MFI of depicted markers in OT-I T_EFF_. **F**. Experimental layout of adoptive transfer experiment. **G**. Lm-OVA CFU in spleen, MLN and PP after adoptive transfer of no OT-I or transfer of splenic, mandLN and MLN T_EFF_. Numbers indicate percentage of organs with Lm-OVA CFU above limit of detection (dotted line). The dotted square highlights OT-I^MLN^-mediated clearance of bacteria in MLN and PP. Data in B are pooled from 2 independent experiments and analyzed using an unpaired t-test. Data in E are from one of two independent experiments and analyzed using ANOVA. Data in G are pooled from 4 independent experiments with each 4-5 mice/group and analyzed using a Kruskal Wallis test against “no OT-I”. *, p < 0.05; **, p < 0.01; ***, p < 0.001.

### mandLN T_EFF_ lack gut-homing receptors but support systemic protection

A hallmark of intestinal mucosa-surveilling inductive lymphoid tissue is the RA-mediated induction of gut-homing trafficking molecules on activated T cells, which promote their subsequent accumulation in lamina propria, GALT and gLN (Mora et al., 2003). To analyze whether this is also the case for mandLN draining the oral mucosa, we transferred OT-I T cells into mice that were subsequently challenged with i.g., i.o. or i.v. Lm-OVA in presence of FTY720 (**Fig. 4C**). All administration routes led to efficient activation of OT-I T cells by day 5 p.i. as assessed by CD44 upregulation (**Fig. 4D and 4E**). After i.g. infection, OT-I T_EFF_ isolated from MLN expressed higher levels of the MAdCAM-1 ligand α4β7 as compared to spleen OT-I T_EFF_ after i.v. infection (**Fig. 4D and 4E**), in line with published observations (Mora et al., 2003; Sheridan et al., 2014). Similarly, CCR9 expression was efficiently induced in MLN T_EFF_ (29.8 ± 6.8% mean ± SEM from 2 independent experiments with n = 8 mice). In contrast, mandLN T_EFF_ failed to increase α4β7 and CCR9 expression (**Fig. 4D and 4E**).

We set out to correlate these data with the protective capacity of mandLN-generated T_EFF_ as compared to those generated in spleen and MLN. We adoptively transferred OT-I CD8^+^ T cells and infected recipient mice separately by i.o., i.v. or i.g. Lm-OVA administration in the presence of FTY720. On day 5 p.i., we isolated OT-I T_EFF_ from mandLN, spleen and MLN, respectively, and transferred equal numbers of effector cells separately into secondary recipient mice, which were i.g. infected with Lm-OVA (**Fig. 4F**). We chose the i.g. route of infection for secondary recipients to induce Lm spread to MLN and PP, as i.v. or i.o. administration of Lm does not result in efficient MLN or intestinal infection (Kursar et al., 2002). This approach therefore allows to assess the potential for T_EFF_ protection in mucosal *versus* systemic sites. Irrespective of the priming site, OT-I T_EFF_ isolated from spleen, MLN and mandLN showed a comparable reduction of bacterial burden in spleen on d 3 p.i. (**Fig. 4G**), while liver was not strongly infected in our setting (not shown). In contrast, mandLN OT-I T_EFF_ did not confer protection to MLN or PP, similar to results obtained after transfer of splenic OT-I T_EFF_ (**Fig. 4G**). In turn, MLN OT-I T_EFF_ caused a reduction in MLN and PP Lm burden in secondary recipients (**Fig. 4G**). This decrease became significant for bacterial loads in MLN when compared to no OT-I transfer or transfer of mandLN and spleen OT-I T_EFF_ in side-by-side comparisons (p < 0.05, Mann-Whitney). In sum, in our model OT-I T_EFF_ generated in oral mucosal-draining mandLN constitute a large fraction of the peripheral T_EFF_ pool early after pathogen ingestion. Our findings further indicate that mandLN T_EFF_ are capable to reduce systemic Lm burden but do not contribute to substantial protection of intestinal lymphoid tissue.

### mandLN stromal cells and DC display low expression of RA-producing enzymes

The lymphoid microenvironment including stromal cells and DC provides critical cues for tissue-selective imprinting of effector homing potential. To characterize the gene expression profile of stromal cells, we performed a single cell RNA sequencing (scRNAseq) analysis of the CD45^-^ TER-119^-^ stromal compartment of mandLN and compared it with stromal cells isolated from MLN and skin-draining inguinal, axillary and brachial peripheral LN (PLN). Unsupervised clustering of combined stromal scRNAseq data identified multiple Col1a1^+^ and Col1a2^+^ fibroblast-like and Cdh5^+^ vascular cell populations (**Fig. 5A and Fig. S3**). While scRNAseq data suggested comparable expression of the RA-producing enzyme Aldh1a1 in fibroblast-like cells of all three LN, expression of Aldh1a2 and Aldh1a3 was higher in MLN than in PLN or mandLN fibroblasts (**Fig. 5B and 5C**). Furthermore, the transcription factor WT1, which drives expression of Aldh1a1 and Aldh1a2 (Buechler et al., 2019), was mainly expressed in MLN fibroblast-like cells (**Fig. 5B and 5C**). To corroborate these data, we performed qPCR analysis for RA-generating enzymes on sorted CD45^-^ TER-119^-^ stromal cells isolated from MLN, PLN and mandLN. This analysis showed that mRNA levels of all three Aldh1a isoforms were higher in MLN as compared to PLN and mandLN stromal cells (**Fig. 5D**). Since DC contribute to gut-homing phenotype imprinting in activated T cells (Erkelens and Mebius, 2017), we isolated CD45^+^ CD11c^+^ MHC-II^high^ DC from MLN, PLN and mandLN and compared mRNA levels of Aldh1a1-3 by qPCR. While Aldh1a1 and Aldh1a3 were expressed at low levels in all CD11c^+^ populations, Aldh1a2 mRNA levels were highest in MLN CD11c^+^ cells (**Fig. 5D**). Taken together, the mandLN microenvironment shares with skin-draining PLN the lack of gut-homing imprinting capacity.

**Figure 5.**
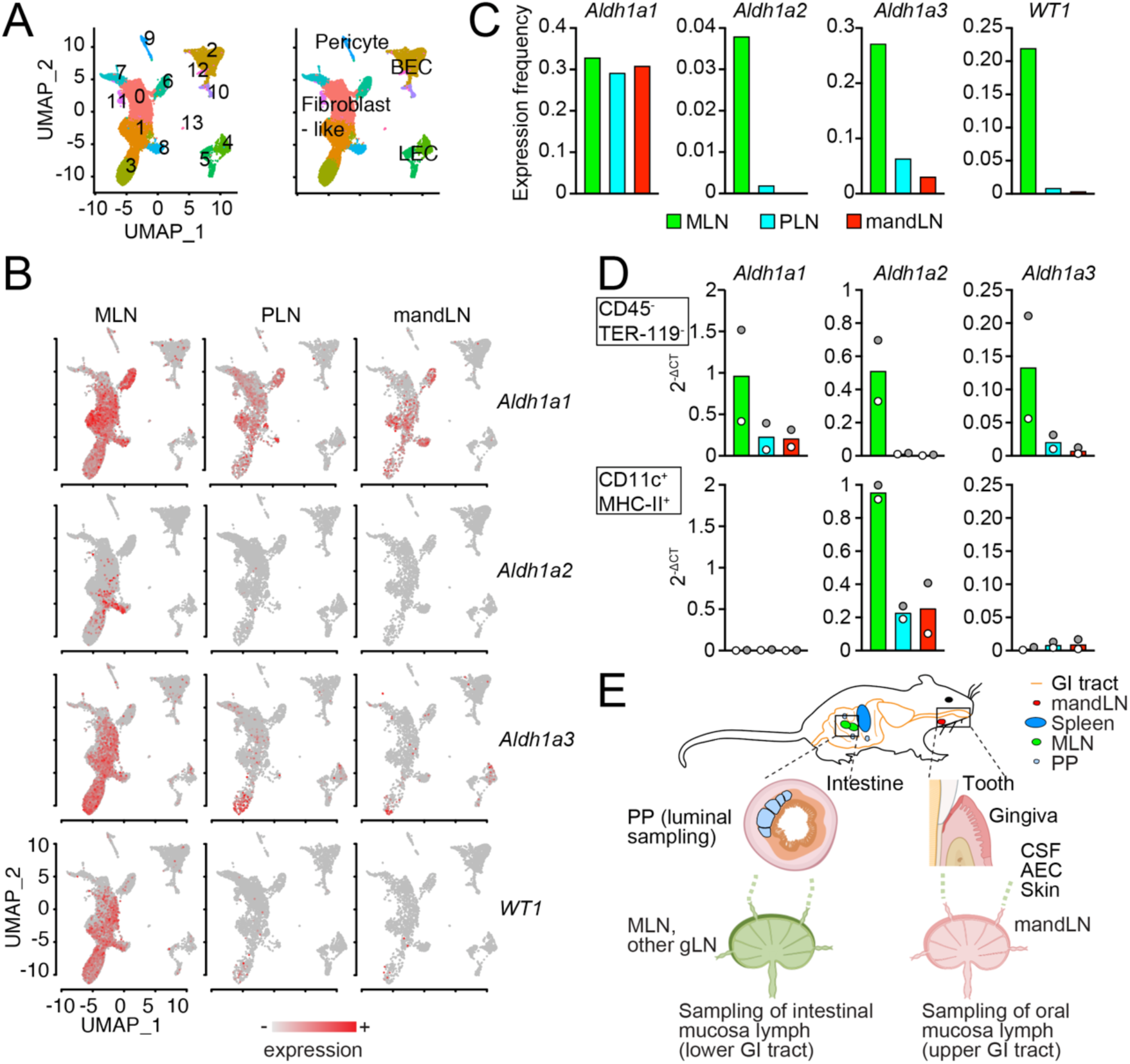
The mandLN microenvironment shows low expression of genes for gut-homing imprinting of effector CD8^+^ T cells. **A**. UMAP clustering of pooled CD45^-^ stromal compartments of MLN, PLN and mandLN based on scRNAseq data. Numbers indicate individual clusters. **B**. Expression of *Aldh1a1, Aldh1a2, Aldh1a3* and *WT1* in MLN (10058 cells), PLN (7750 cells) and mandLN stroma (4434 cells) based on scRNAseq data. **C**. Expression frequency of *Aldh1a1, Aldh1a2, Aldh1a3* and *WT1* in fibroblast-like cells based on data in B. **D**. *Aldh1a1, Aldh1a2, Aldh1a3* expression in CD45^-^ TER-119^-^ stromal cells and CD11c^+^ MHC-II^+^ DC isolated from of MLN, PLN and mandLN assessed by qPCR. Shown are the 2^-ΔCT^ means of duplicates from two independent experiments (grey and white fill). Bars represent mean. **E**. Graphical summary. CSF, cerebrospinal fluid; AEC, anterior eye chamber.

The mucosa lining the oral cavity is the first site of contact with ingested microbes before their passage to the esophagus, stomach and intestinal tract. The presence of lymphatic vessels in the mucosa lining the oral cavity suggests a continuous surveillance of regional lymph by sentinel LN (Ando et al., 2011; Ushijima et al., 2008). The sequestration of orally administered Lm in mandLN observed here suggests that lymphatic drainage leads to continuous sampling of the oral microbiome at this anatomical localization, confirming observations made after oral delivery of *Trypanosoma cruzi* (Barreto de Albuquerque et al., 2018; Silva dos Santos et al., 2017). It remains unclear how Lm entry into oral lymphatic vessels is regulated. In the oral cavity, the gingival sulcus, which is the space between the gingiva and teeth, is a particularly vulnerable site exposed to trauma caused by mastication and biting. Its epithelium is non-keratinized and transitions to the junctional epithelium that binds directly to teeth (Moutsopoulos and Konkel, 2018; Gaffen and Moutsopoulos, 2020). Conceivably, lymphatic vessels below the crevicular epithelium lining the gingival sinus may participate in collecting microbe-containing tissue fluids for transport to mandLN. In addition, the vascular-rich sublingual mucosa has absorptive properties, which is clinically relevant to systemically administer drugs or vaccines (Gaffen and Moutsopoulos, 2020). Oral microbiota might be collected here for lymphatic transport to regional LN, a process which may be further facilitated by the lack of a thick mucus layer as is present in intestinal mucosa. These observations do not exclude a role for migratory DC in transporting Ag from the oral mucosa to mandLN. Furthermore, our study does not address how the presence of food, on which Lm is usually found, affects microbial drainage to mandLN or its passage through the stomach and small intestine. It has been reported that infection with food-borne Lm leads to the appearance of endogenous T_EFF_ in MLN and PP one week post Lm infection (Sheridan et al., 2014). While mandLN had not been analyzed in that study, these data are consistent with the kinetics of Lm dissemination to MLN and PP reported here.

One of the pillars of cellular immune responses is the imprinting of tissue-specific homing molecules during T cell activation. In skin-draining LN, sunlight-generated vitamin D3 imprints expression of CCR4, CCR10 and P- and E-selectin ligands, while GALT and gLN process dietary carotenes and retinol to imprint a gut-homing *α*_4_ *β*_7_^+^ CCR9^+^ phenotype (Iwata et al., 2004; Sigmundsdottir and Butcher, 2008). This remarkable “division of labor” ensures an optimal use of resources to direct T cell responses to the anatomical site of pathogen entry. Most of this paradigm has been established by analyzing lymphocyte activation in lymphoid tissue of the lower GI tract. This raises the question why mandLN differ from those sites, even though both drain lymph from microbe-rich mucosal tissue. One explanation is that the induction of RA-generating enzymes in migratory CD103^+^ DC for gut-homing phenotype imprinting in T cells requires RA generated by epithelial cells of the small intestine (Molenaar et al., 2011; Johansson-Lindbom et al., 2005; Larange and Cheroutre, 2016). Similarly, we observed low expression of RA-producing enzymes in stromal cells of mandLN, suggesting that RA is not present in sufficiently high concentration in oral cavity-draining mandLN to direct substantial T_EFF_ trafficking towards gut. In this context, mandLN do not only drain lymph from the oral mucosa but also of the anterior eye chamber, NALT and skin as well as cerebrospinal fluid of the central nervous system (CNS) (Boonman et al., 2004; Pabst, 2015; Ma et al., 2019; Van den Broeck et al., 2006) (**Fig. 5E**). This may further influence the expression of genes that direct homing patterns in dendritic and stromal cells, e.g. to minimize loss of CD8^+^ T_EFF_ to intestinal sites during CNS inflammation. In line with our findings, i.n. Lm infection leading to Ag presentation in NALT does not confer intestinal immunity (Sheridan et al., 2014).

An open question is to which extent mandLN priming contributes to the generation of systemic and oral mucosa CD8^+^ memory cells. After adoptive transfer of mandLN OT-I T_EFF_ into infection-matched recipients, we recovered memory T cells in spleen but not in MLN or PP (not shown). Furthermore, the requirements for entry into the oral cavity are not well defined to date and we found lymphocytes difficult to extract from this location in a quantitative manner (not shown). Of note, 2PM imaging identified memory T cells in the oral mucosa at > 30 d post i.o. Lm infection (**Video S4**), suggesting that mandLN may contribute to local memory generation of their surveilled barrier tissue.

PP and gLN constitute a complementary surveillance system of the lower GI tract, with PP containing gut lumen-sampling M-cells and gLN draining lymphatic vessels originating in lamina propria. Our data suggest that mandLN form together with M-cell-containing MALT including tonsils (in humans) and NALT a comparable “dual surveillance” system for the oropharyngeal section of the upper GI tract (**Fig. 5E**). The role for mandLN in this process has thus far remained largely overlooked, since most immunologists use i.v. infection of Lm as a robust systemic infection model or bypass the oral cavity by i.g. infection. Taken together, our study adds the increasingly acknowledged site-specific imprinting of host immune responses along the length of the GI tract, consistent with an instructional role for the Ag sampling location (Mowat and Agace, 2014; Esterhazy et al., 2019).

## Material and methods

### Mice

Female C57BL/6JRj (Janvier, Le Genest-Saint-Isle, France), CCR7^-/-^ (Forster et al., 1999), Tg(Itgax-Venus)1Mnz “CD11c-YFP” (Lindquist et al., 2004) and Tg(Prox1-EGFP)KY221Gsat “Prox1-GFP” (Choi et al., 2011) mice were used for imaging, as recipients for T cell adoptive transfer or Lm infection. Polyclonal T cells were isolated from C57BL/6 or “Ubi-GFP” donors (Schaefer et al., 2001). Tg(TcraTcrb)1100Mjb OT-I TCR mice (Hogquist et al., 1994) backcrossed on a GFP^+^ (Schaefer et al., 2001), dsRed^+^ (Kirby et al., 2009) or tdT^+^ (Madisen et al., 2010; de Vries et al., 2000) background were described before (Ficht et al., 2019). In some experiments, CD45.1^+/+^ or CD45.1^+^/CD45.2^+^ OT-I were used. All animals were bred in specific pathogen-free conditions at the Central animal facility of the University of Bern, and University of Fribourg. All animal work has been approved by the Cantonal Committees for Animal Experimentation and conducted according to federal guidelines.

### T cell purification and adoptive transfer

Spleens and LN were dissociated using 70 μm cells strainers, and CD8^+^ T cells were negatively isolated using the EasySep^™^ Mouse CD8^+^ T cell Isolation Kit (Stem Cell Technologies, Grenoble, France) or MojoSort^™^ Isolation Kits (BioLegend, San Diego, US) according to the manufacturer’s protocol. Purity of isolated CD8^+^ T cells was > 90%. For 2PM imaging, polyclonal CD8^+^ T cells were labelled with 20 μM CellTracker^™^Blue CMAC (7-amino-4-chloromethylcoumarin; Invitrogen) for 20-30 min at 37° C. For proliferation assays, OT-I T cells were labelled with 5 μM CellTrace^™^ Violet (Invitrogen) at 37° C for 20-30 min.

### Bacterial infection and CFU quantification

Lm strain 10403s expressing ovalbumin (Lm-OVA) (Zehn et al., 2009) or GFP (Lm-GFP) (Abrams et al., 2020) and *Lm* strain EGD-e, carrying a recombinant InlA with S192N and Y369S mutations (Lm-InlA^m^) (Monk et al., 2010) were kindly provided by Profs. Doron Merkler (University of Geneva, Switzerland), Neal M. Alto (University of Texas, Southwestern Medical Center, US) and Colin Hill (University College Cork, Ireland), respectively. Bacteria from glycerol stocks were grown in Brain heart infusion (BHI) medium until mid-log phase was reached. Prior and after infection, mice were deprived of food and water for 4 h and at least 15 min respectively, and received 2 ⨯ 10^9^ CFU in the oral cavity by pippeting into the mouth over 1-5 min as described (Barreto de Albuquerque et al., 2015) or directly into the stomach by intragastric gavage, or i.v. 5 ⨯ 10^3^ CFU. For Lm titration, 10^9^, 10^8^, 10^7^ and 10^6^ CFU were used. Bacterial suspensions were serially diluted and plated on BHI agar plates to verify the actual number of CFU in the inoculum.

For CFU quantification, mandLN, MLN, PP and spleen were aseptically harvested, homogenized in a 70 µm nylon filter and lyzed in sterile water. Alternatively, organs were homogenized using MagNALyser tubes Green Beads (Roche, France) containing sterile PBS (5000 rpm for 1 min). Serial dilutions of homogenates were plated on BHI agar, and colonies were counted after 24 h of incubation at 37°C. For PP, BHI agar plates contained 200 μg/mL streptomycin to prevent gut bacteria contamination, since the 10403s strain is strepromycin resistant. CFU was adjusted according to the dilutions and calculated per number of LN or PP collected. For protection experiments, spleen, liver, PP and MLN were lysed in 2-mL tubes containing one steal bead and 1 mL PBS + 0.1% Tween20 (Sigma-Aldrich, St. Louis, USA), using a QiaTissueLyzer (Qiagen, Venlo, Netherlands) at 25 Hz, 3 min. Serial dilutions were plated on BHI agar, and colonies were counted after 48 h of incubation at 30°C. For PP, BHI agar plates contained 200 μg/mL streptomycin. CFU was adjusted according to the dilutions and calculated per gram of organ.

### FTY720 treatment

Lymphocyte egress from lymphoid tissue was blocked by treating mice intraperitoneally with 2 mg/kg FTY720 (Sigma), a sphingosine-1-phosphate receptor 1 (S1PR1) inhibitor, starting at d 1 p.i. with repeated doses after 48 h.

### Flow cytometry

LN, PP and spleen were harvested at the indicated time points and single cell suspensions were obtained by passing organs through 70 μm cell strainers. Red blood cell lysis was performed on splenocytes. Counting of viable cells negative for Trypan blue was performed using Neubauer chamber. Fc receptors were blocked with purified anti-CD16/CD32 mAb (2.4G2) in FACS buffer (PBS with 2% FCS, 2 mM EDTA and 0.05% NaN_3_) for 10 min. Cell surface stainings were performed with the appropriate combinations of mAbs on ice for 30 min in the dark and washed in FACS buffer (**Table 1**). Zombie Aqua^™^, Zombie Violet^™^ or Zombie Red^™^ (BioLegend) were used to discriminate live and dead cells. After staining, cells were fixed for 20 min with 2% paraformaldehyde, washed and resuspended in PBS. Acquisition was performed using LSR Fortessa (BD Bioscience), LSR II (BD Bioscience), LSR II (BD Bioscience) or Attune NxT Flow cytometer (Thermo Fisher) and analyzed using FlowJo (TreeStar) software.

**Table 1.**

| Antibody/streptavidin | Fluorochrome | Clone | Concentration<br>(µg/mL) | Company | Reference<br>number |
| --- | --- | --- | --- | --- | --- |
| CD103 | APC | 2 E7 | 2 | BioLegend | 121414 |
| CD11b | PE-Cy7 | M1/70 | 0.4 | BioLegend | 101215 |
| CD11c | BV605 | N418 | 2 | BioLegend | 117334 |
| CD127 | APC | A7R34 | 2 | BioLegend | 135012 |
| CD19 | biotinylated | 6D5 | 5 | BioLegend | 115503 |
| CD199 (CCR9) | PE-Cy7 | CW1.2 | 1 | BioLegend | 128711 |
| CD25 | APC | PC61 | 1 | BioLegend | 102012 |
| CD44 | biotinylated | IM7 | 2.5 | BD<br>Biosciences | 553132 |
| CD44 | PerCP | IM7 | 2 | BioLegend | 103036 |
| CD44 | PerCP-Cy5.5 | IM7 | 2 | BioLegend | 103032 |
| CD45.1 | PE | A20 | 1 | BioLegend | 110708 |
| CD45.2 | AF488 | A20 | 2.5 | BioLegend | 110718 |
| CD62L | BV421 | MEL-14 | 0.5 | BioLegend | 104436 |
| CD69 | BV711 | H1.2F3 | 1 | BioLegend | 104537 |
| CD69 | PE | H1.2F3 | 1 | BioLegend | 104508 |
| CD8 | APC/Fire 750 | 53-6.7 | 1 | BioLegend | 100766 |
| CD80 | FITC | 16-10A1 | 2.5 | BioLegend | 104705 |
| CD86 | APC-Fire 750 | GL-1 | 1 | BioLegend | 105045 |
| CXCR3 | BV510 | CXCR3-173 | 2 | BioLegend | 126527 |
| CXCR3 | BV605 | CXCR3-173 | 2 | BioLegend | 126523 |
| I-A/I-E (MHCII) | BV421 | M5/114.15.2 | 0.5 | BioLegend | 107632 |
| KLRG1 | PE-Cy7 | 2F1 | 1 | BioLegend | 138415 |
| LPAM-1 ($\alpha_4\beta_7$) | APC | DATK32 | 2 | eBioscience | 17-5887-82 |
| Ly-6G/C | biotinylated | Gr-1 | 2.5 | BioLegend | 108403 |
| NK1.1 | PE | PK136 | 1 | BioLegend | 108707 |
| streptavidin | BV711 | – | 0.5 | BioLegend | 405241 |
| streptavidin | PE | – | 2 | BioLegend | 405203 |
| TCRβ | biotinylated | H57-597 | 2.5 | BioLegend | 109203 |

For flow cytometry analysis of DCs, LN were cut in small fragments and digested in type IV collagenase (1 mg/mL, Worthington) and DNase I (100 U/mL; Roche) in 24-well plates under rotation at 20 rpm for 15-20 min at 37°C. Tissue fragments were dissociated by pipetting using a Pasteur pipette and incubated for additional 15 min. In case tissue fragments were not completely dissociated by pipetting using a 1 mL microtiter pipette, samples were incubated for additional 10 min. For sorting of DC and stromal cells, we used collagenase P (0.5 mg/mL; Roche), Dispase II (0.5 mg/mL; Roche) and DNase I (100 U/ml; Roche) for digestion in a 5-mL round bottom tube.

### Intravital imaging of mandLN and gingiva

Mice were anesthetized by i.p. injection of 8-10 µL/g Ketamine (20 mg/mL) and xylazine (1 mg/mL) and after 15 min, 30 µL i.p. of acepromazine (2.5 mg/mL).

Anesthesia was supplemented when needed with ketamine/xylazine (half dose of xylazine). The fur was removed from the operating area using an electric razor followed by hair removal cream. Mice were placed on a customized stage used for submandibular salivary gland (SMG) imaging (Ficht et al., 2018). Neck and teeth were fixed to reduce shifting. To expose mandLNs, a 0.5 - 1 cm incision was made along the neck region in the skin above the left lobe of the SMG. Under the stereomicroscope, surrounding connective tissue between left and right lobes of the SMG, and between the left lobe and the adjacent skin were disrupted to set the left lobe free. Blood and lymphatic vessels, and the mandLN were kept untouched and moisturized with saline during the surgical procedure. The left lobe of the SMG was pulled on top of a coverslip on a holder and glued using veterinary adhesive (Vetbond, M3). A ring of grease (Glisseal N, VWR) was made in order to create space to be filled with saline and keep the tissues moisturized. The SMG was fixed in a position to expose the mandLN and fat tissue was carefully removed. Another holder with coverslip was placed on top and touching the LN without impairing the blood flow. To maintain physiological temperature, a heating ring was connected to a water bath.

To expose the gingiva, a 0.5 cm incision was made in the lower lip using an electric cauterizer. Sutures were tied to both sides and fixed into the surface of the surgical stage to stretch the area and expose the inferior teeth. Viscotears (Alcon) was used to keep the tissue moisturized and a holder with a coverslip was placed on top. To maintain physiological temperature, a heating ring was connected to a water bath and a temperature probe was used. For 2PM imaging of Lm accumulation in mandLN, we pipetted 20 µL Lm-GFP (4-8 x 10^8^ CFU/ml) into the oral cavity and surgically prepared mice 1-2 h later by exposing gingiva and mandLN as described above. After addition of another 10 μL Lm-GFP to gingiva, we analyzed mandLN by 2PM imaging.

2PM was performed with an Olympus BX50WI microscope equipped with 20X Olympus (NA 0.95) or 25X Nikon (NA 1.0) objectives and a TrimScope 2PM system controlled by ImSpector software (LaVisionBiotec). Prior to recording, Alexa Fluor 633- or Alexa Fluor 488 conjugated MECA-79 (10 μg/mouse) was injected i.v. to label high endothelial venules (HEV). For 2-photon excitation, a Ti:sapphire laser (Mai Tai HP) was tuned to 780 or 840 nm. For 4-dimensional analysis of cell migration, 11 to 16 x-y sections with z-spacing of 4 μm were acquired every 20 s for 20-30 min; the field of view was 150-350 x 150-350 μm. Emitted light and second harmonic generation (SHG) signals were detected through 447/55 nm, 525/50 nm, 593/40 nm and 655/40 nm bandpass filters with non-scanned detectors when using C57BL/6 recipient mice. For CD11c-YFP^+^ recipient mice, 447/55 nm, 513/20 nm, 543/39 nm and 624/30 nm were used as bandpass filters. SHG or HEV signal were used as anatomical reference channel for real-time offset correction to minimize tissue shift (Vladymyrov et al., 2016). Sequences of image stacks were transformed into volume-rendered four-dimensional videos with using Imaris software (Bitplane), which was also used for semi-automated tracking of cell motility in three dimensions. Cell centroid data was used to calculate key parameters of cell motility using Matlab (R2019b, MathWorks). Speed was defined as total track length divided by total track duration in µm/min. The arrest coefficient was derived from the percentage of time a cell is migrating below a motility threshold speed of 5 µm/min. Meandering index was calculated by dividing displacement by track length. Since the meandering index is influenced by track duration (Beltman et al., 2009), we calculated the corrected track straightness defined as meandering index multiplied by the square root of cell track duration.

### Stereomicroscope imaging

MandLN and gingiva were exposed and pictures were taken using a Leica MZ16 FA stereomicroscope equipped with color high resolution camera (Leica). Images were processed using Adobe Photoshop CS6.

### Immunofluorescence of LN sections

Mice were anesthetized with i.p. injection of ketamine and xylazine and perfused with cold 2% PFA. MandLN were harvested and fixed overnight in 4% PFA, and dehydrated in 30% sucrose overnight prior to embedding in TissueTek O.C.T. compound (Sakura) for cryostat sectioning. Slides with 10 μm-thick cryosections were mounted with ProLong^™^ Gold Antifade Mountant (Molecular Probes). Fluorescence microscopy was performed using Leica SP5 confocal microscope with 20X (NA 0.7) and 63X (NA 1.3) Leica objectives. Images were processed using Adobe Photoshop CS6 and Imaris 8.4.1 (Bitplane). Brightness and contrast were adjusted for each image individually.

### Single cell gene expression and data analysis

Single cell gene expression of CD45^-^ stromal cells was measured using the 10x Chromium system, with the Next GEM Single Cell 3’ Reagent Kit v3.1 (10x Genomics, Pleasanton, CA, USA). GEM generation and barcoding, reverse transcription, cDNA amplification and 3’ Gene Expression library generation steps were all performed according to the manufacturer’s user guide. Specifically, the designated volume of each cell suspension (800 -1200 cells/µL) and nuclease-free water were used for a targeted cell recovery of 10’000 cells according to the Cell Suspension Volume Calculator Table of the abovementioned user guide. GEM generation was followed by a GEM-reverse transcription incubation, a clean-up step and 15 cycles of cDNA amplification. The quality and quantity of the cDNA was assessed using fluorometry and capillary electrophoresis, respectively. The barcoded cDNA libraries were pooled and sequenced paired-end and single indexed on an Illumina NovaSeq 6000 sequencer using a S2 flowcell (100 cycles). The read setup was as follows: read 1: 28 cycles, i7 index: 8 cycles, i5: 0 cycles and read 2: 91 cycles. An average of 740,045,087 reads/library were obtained, which corresponds to an average of 74,091 reads/cell.

Mapping and counting of the UMIs for the samples from mandLN, MLN and PLN were performed using Cellranger (version 3.0.2, 10x Genomics) with the reference genome GRCm38.93 from Ensembl to build the necessary index files. Subsequent analysis was performed in R (version 4.0.2) (R Core Team, 2016). The Scater package (version 1.14) (McCarthy et al., 2017) was used to assess the proportion of ribosomal and mitochondrial genes as well as the number of detected genes. Cells were considered as outliers and filtered out if the value of the proportion of expressed mitochondrial genes or the number detected genes deviated more than three median absolute deviations from the median across all cells. After quality control, the sample from mandLN retained 4434 cells, the sample from MLN retained 10058 cells and the sample from PLN retained 7750 cells. Normalization between samples was done with the deconvolution method of Lun et al. (Lun et al., 2016b) using the package Scran (version 1.14) (Lun et al., 2016a). Samples were integrated with the FindIntegrationAnchors function of the package Seurat (version 3.1) based on the first 20 principal components (PCs) (Stuart et al., 2019). Graph-based clustering was done with the FindNeighbors and FindClusters functions of the Seurat package using the first 40 PCs from the dimensionality reduction step. The Clustree package (version 0.4) (Zappia and Oshlack, 2018) was used to determine the resolution resulting in clustering concurring with the presumed cell types, which was 0.4. Clusters were annotated based on marker genes that were identified with the FindMarkers function of Seurat.

Following this, seven clusters were removed from the analysis as they represented undesired hematopoietic cells. The remaining cells were re-clustered in an identical fashion but with a resolution of 0.3.

### Quantitative PCR

For sorting of DC and stromal cells, cells suspensions were stained with mAbs (**Table 1**) and propidium iodide to discriminate between live *versus* dead cells. CD45^+^ MHC-II^high^ CD11c^+^ DCs and CD45^-^ stromal cells were sorted using a FACSAria Fusion (BD). For real-time quantitative RT-PCR (qPCR), the total RNA from 2.7-30 ⨯ 10^4^ sorted DC and stromal cells was extracted using Trizol Reagent (Sigma-Aldrich) and coprecipitant GlycoBlue^™^ (Invitrogen). Reverse transcriptase reactions were performed using High-Capacity cDNA Reverse Transcription kit (Applied Biosystems, USA) according to the manufacturer’s instructions. Real-time RT-PCR assays were performed on StepOnePlus (Applied Biosystems, USA) using FastStart Universal SYBR Green Master (Rox) (Roche, Switzerland). QuantiNova LNA-enhanced primers for *Aldh1a1, Aldh1a2, Aldh1a3* and as endogenous housekeeping reference gene hypoxantine-guanine phosphoribosyltransferase (HPRT) were purchased from Qiagen (Germany) (**Table 2**). Real-time PCR reactions were performed in duplicates in a total volume of 20 μL. The cycling conditions were: 95°C for 10 min, followed by 40 cycles at 95°C for 10 s and 60°C for 30 s. After amplification, dissociation curves were performed to monitor primers specificity, revealing only one melting peak for each amplified fragments. qPCR data were normalized to the housekeeping gene HPRT and the relative changes in mRNA expression were calculated by 2^-ΔCt^.

**Table 2.**

| Assay Name | Full name | Gene ID | Ref Seq | Catalogue # |
| --- | --- | --- | --- | --- |
| MM_HPRT_2521490 | hypoxanthine guanine phosphoribosyl transferase | 15452 | NM_013556 | SBM1225379 |
| MM_ALDH1A1_2045606 | aldehyde dehydrogenase family 1, subfamily A1 | 11668 | NM_001361506 | SBM0801506 |
| MM_ALDH1A2_1966774 | aldehyde dehydrogenase family 1, subfamily A2 | 19378 | NM_009022 | SBM0722689 |
| MM_ALDH1A3_2125665 | aldehyde dehydrogenase family 1, subfamily A3p | 56847 | NM_053080 | SBM0881559 |

### Statistical analysis

Student’s t-test, Mann-Whitney U-test, ANOVA or Kruskal-Wallis test were used to determine statistical significance as indicated (Prism, GraphPad). Significance was set at p < 0.05.

### Supplemental material

Fig. S1. Depiction of surgery for gingiva and mandLN.

Fig. S2. Flow cytometry of DC and CD8^+^ T cell subsets.

Fig. S3. Expression of selected genes of interests by scRNAseq.

Video S1. OT-I in uninfected mandLN.

Video S2. OT-I in mandLN on d 2 post i.o. Lm infection.

Video S3. OT-I in mandLN on d 3 post i.o. Lm infection.

Video S4. OT-I in gingiva in memory phase.

## Author contribution

JBdA performed experiments with help from DvW, LMA and XF. JA, GvG and DF performed scRNAseq analysis under supervision of RB. XF and MI provided a script for cell tracking analysis. JBdA, CM and JVS designed experiments and wrote the manuscript with input from all coauthors.

## Acknowledgments

This work benefitted from the Microscopy Imaging Center of the University of Bern and the BioImage Light Microscopy Facility and Cell Analytics Facility of the University of Fribourg. We thank Daniela Grand, Antoinette Hayoz and Pablo Bànicles for expert technical assistance, Pamela Nicholson, Catia Coito, Marion Ernst and Daniela Steiner from the Next-Generation Sequencing Platform of the University of Bern and Profs.

Doron Merkler, Neal Alto and Colin Hill for sharing Listeria strains.

## Supplemental Figure legend

**Figure S1.**
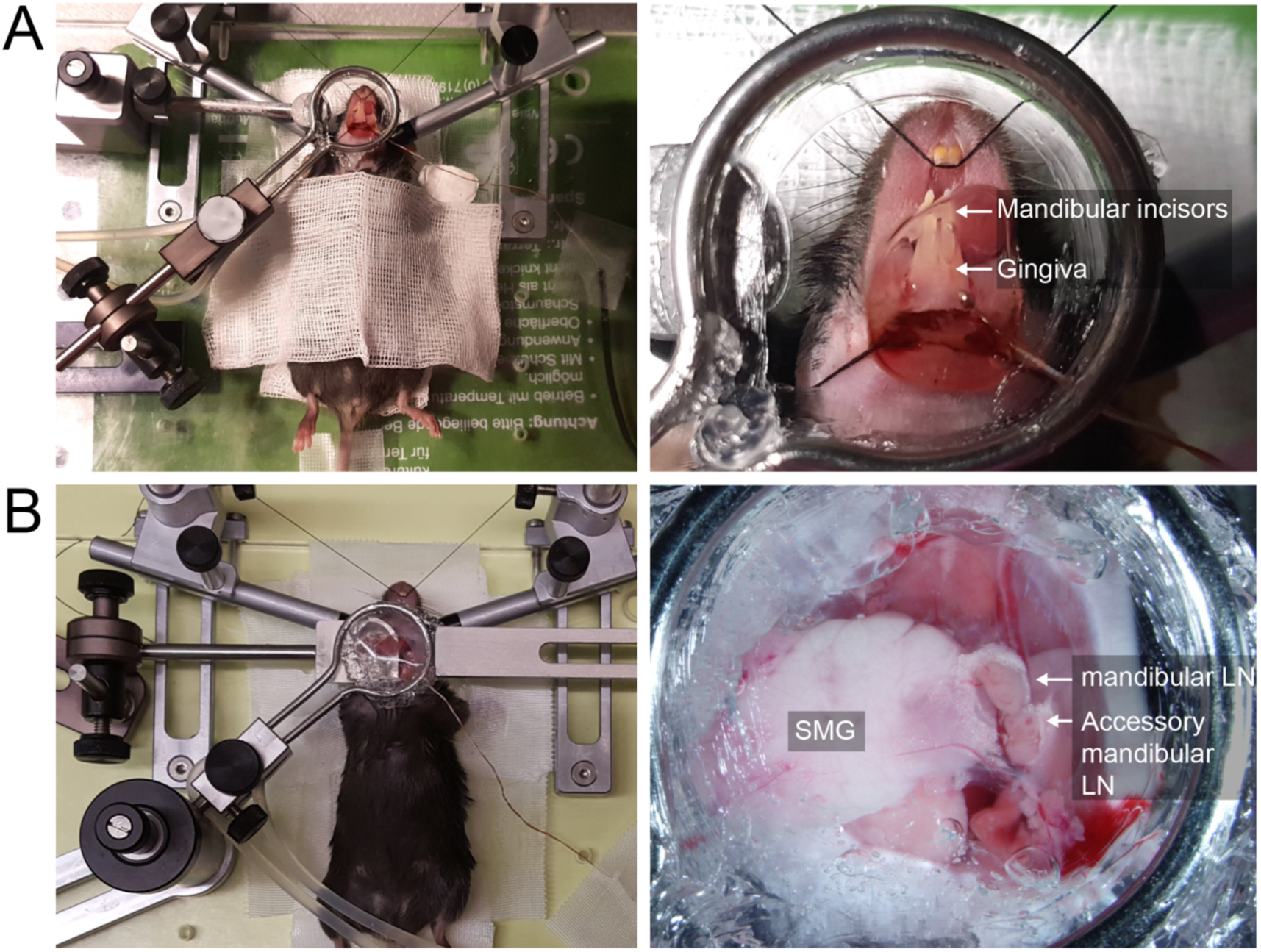
Surgical setup (left panels) and stereomicroscope images (right panels) of gingiva (**A**) and mandLN (**B**). SMG, submandibular salivary gland.

**Figure S2.**
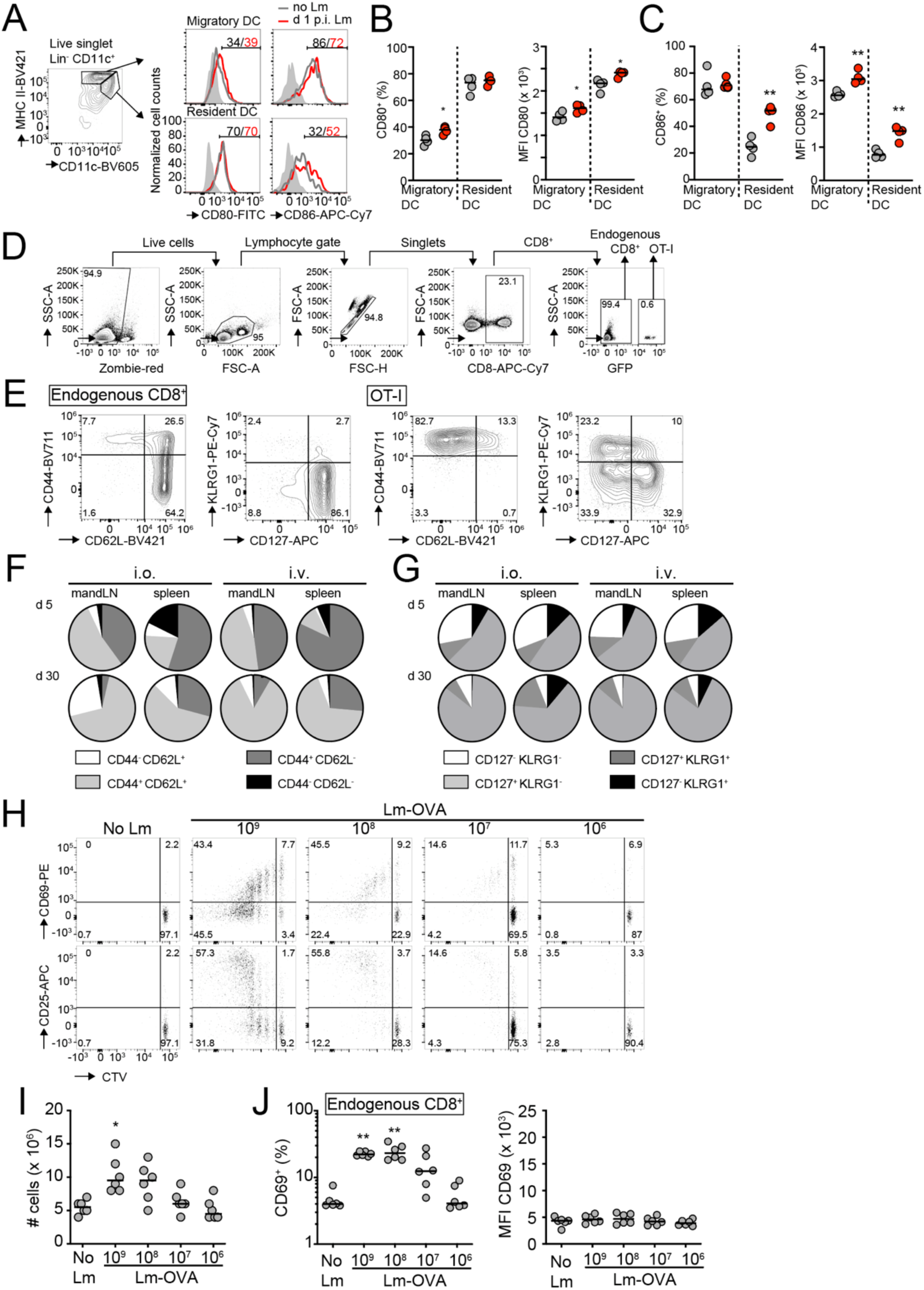
Flow cytometry analysis of DC and CD8^+^ T cells. **A**. Representative flow cytometry plot for migratory and resident DC and CD80/CD86 expression in each DC population. Gray shade, FMO; gray line, no Lm; red line, after i.o. Lm-OVA infection. Numbers indicate percent expressing cells. **B and C**. CD80 (**C**) and CD86 (**D**) expression and MFI on migratory and resident DC. Data in B and C are from one of two independent experiments with 4 mice/group and analyzed using an unpaired t-test. **D and E**. Examples of gating strategy (D) and flow cytometry plots (E). **F and G**. Pie charts showing the proportion of OT-I populations based on CD44/CD62L (**F**) and KLRG1/CD127 (**G**) expression on d 5 and 30 following i.o. and i.v. infections. **H**. Representative flow cytometry plots of CD69/CD25 expression and proliferation in mandLN OT-I T cells on d 3 p.i. with decreasing Lm-OVA inoculum. **I**. MandLN cellularity on d 3 after i.o. Lm-OVA infection. **J**. CD69 expression and MFI CD69 on endogenous mandLN CD8^+^ cells with decreasing Lm-OVA inoculum. *, p < 0.05; **, p < 0.01.

**Figure S3.**
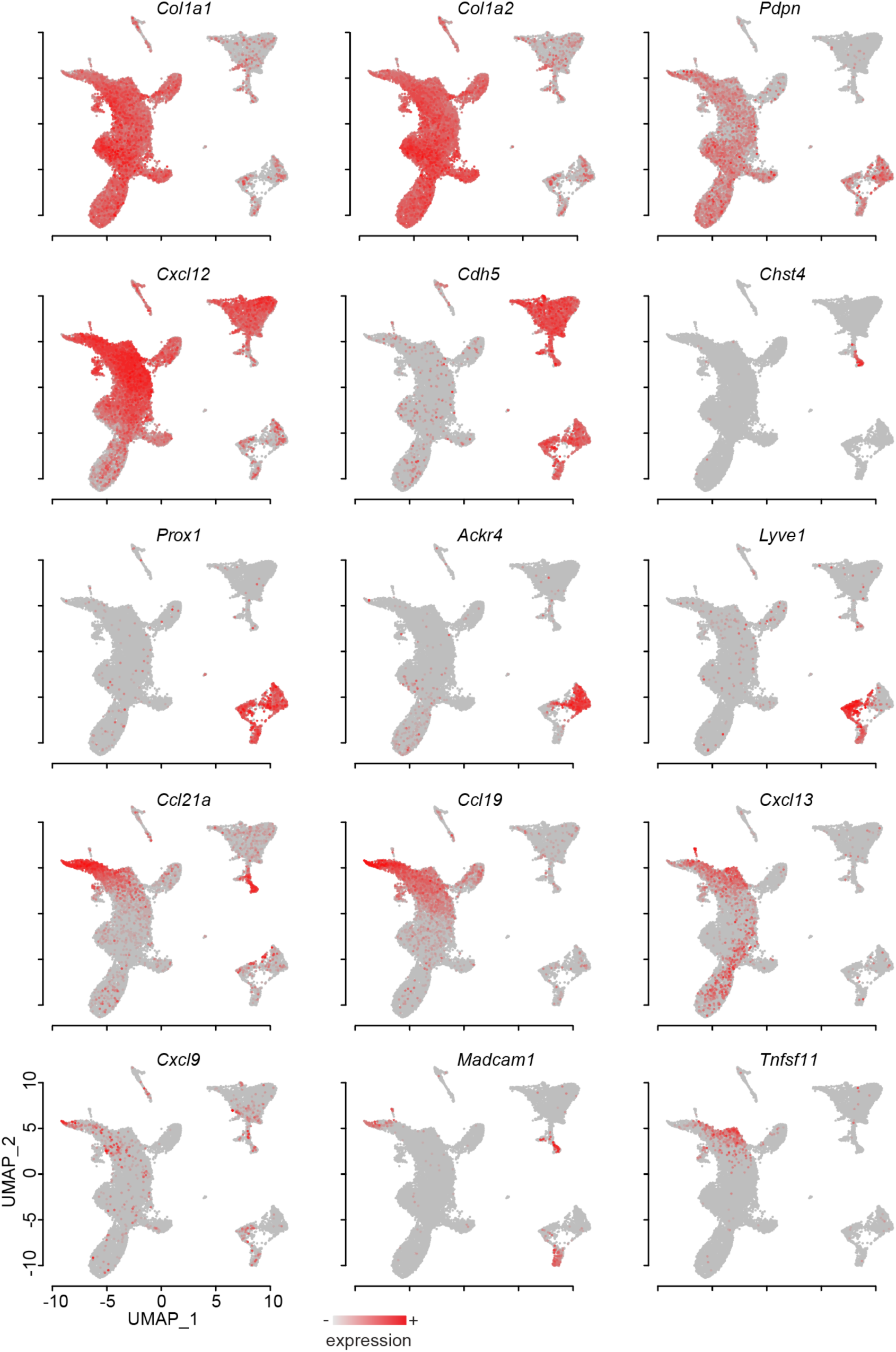
Expression of selected genes of interests in stromal cell clusters by scRNAseq. *Col1a1*, collagen type I alpha 1 chain; *Col1a2*, collagen type I alpha 2 chain; *Pdpn*, podoplanin; *Cdh5*, VE-cadherin; *Chst4*, Carbohydrate Sulfotransferase 4; *Tnfsf11*, TNF Superfamily Member 11 (RANKL).

## Supplementary video legend

**Video S1**. OT-I motility in mandLN in the absence of infection. Scale bar, 30 µm; time in min:s.

**Video S2**. Polyclonal CD8^+^ and OT-I T cell motility in mandLN on d 2 post i.o. Lm-OVA infection. Scale bar, 30 µm; time in min:s.

**Video S3**. Polyclonal CD8^+^ and OT-I T cell motility in mandLN on d 3 post i.o. Lm-OVA infection. Scale bar, 30 µm; time in min:s.

**Video S4**. OT-I T cell migration in gingiva in memory phase following i.o. Lm-OVA infection. Scale bar, 50 µm; time in min:s.

